# Bioaccumulation of heavy metals in commercially important fish species from the tropical river estuary suggests higher potential child health risk than adults

**DOI:** 10.1101/681478

**Authors:** A. S. Shafiuddin Ahmed, Sharmin Sultana, Ahasan Habib, Hadayet Ullah, Najiah Musa, Md. Mahfujur Rahman, Md. Shafiqul Islam Sarker

## Abstract

The Karnaphuli, a major river of Bangladesh, located off the coast of Chittagong in the Bay of Bengal is largely exposed to the heavy metal pollutants, which may be toxic to humans and aquatic fauna. The estuary is a striking example of a site where human pressure and ecological values collide with each other. In spite of being a major supplier of fish food for local community, there has been no study carried out to date to assess the potential human health risk due to heavy metal contamination in the fish species from this estuary. Therefore, the aim of present study was to assess bioaccumulation status and the potential human health risk evaluation for local consumers. Six commercially important fish species, *Apocryptes bato, Pampus chinensis, Hyporhamphus limbatus, Liza parsia, Mugil cephalus, and Tenualosa toil* from the Karnaphuli River estuary were collected to analyze heavy metals concentration level. Heavy metals As, Pb, Cd, Cr and Cu were detected from the samples using inductively coupled plasma mass spectrometry (Model: ELAN9000, Perkin-Elmer, Germany). The hierarchy of the measured concentration level of the metals was as follows: Pb (mean: 13.88, range: 3.19 - 6.19) > Cu (mean: 12.10, range: 10.27 - 16.41) > As (mean: 4.89, range: 3.19 – 6.19) > Cr (mean: 3.36, range: 2.46 – 4.17) > Cd (mean: 0.39, range: 0.21 - 0.74). The Fulton’s condition factor denoted that organisms were particularly in better ‘condition’ and most of the species were in positive allometric growth. The Bioaccumulation factors (BAFs) observed in the species of the contaminants were organized in the following ranks: Cu (1971.42) > As (1042.93) > Pb (913.66) > Cr (864.99) > Cd (252.03), and among all the specimens, demersal fish, *A. bato* corresponded to the maximum bio-accumulative organism. Estimated daily intake (EDI), target hazard quotient (THQ) and carcinogenic risk (CR) assessed for human health risk implications suggest that the values are within the acceptable threshold for all sorts of consumers. Hence none of them would experience non-carcinogenic and carcinogenic health effect for the ingestion of the fishes. However, children are shown to be largely susceptible than adults to non-carcinogenic and carcinogenic health effect due to the consumption of fish. Therefore, an appropriate guidlines and robust management measures needed to be taken to restore the estuarine health condition for greater benefit of the quality of fish products for local consumption.

**Graphical abstract:** 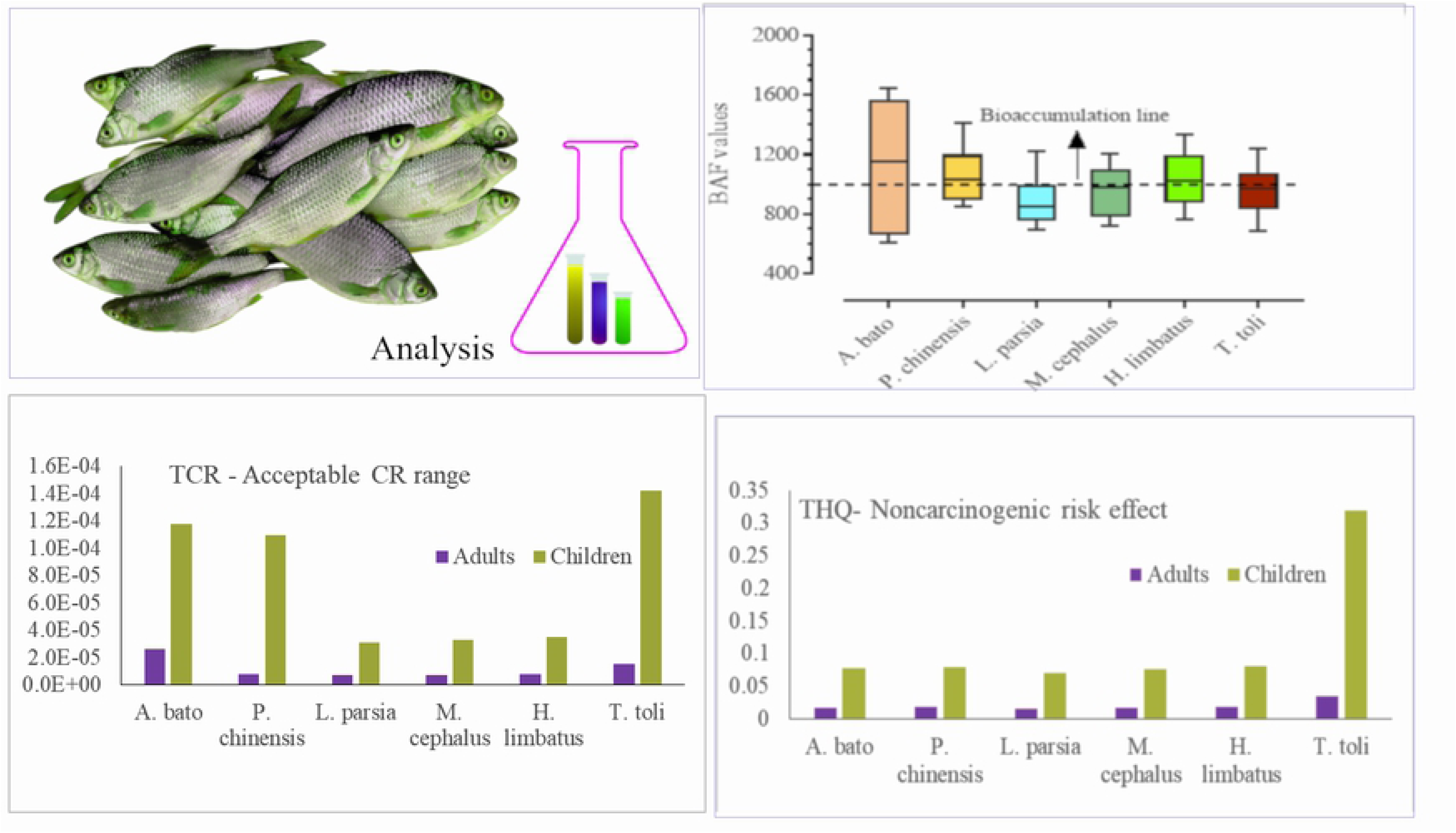

## Introduction

All living organisms including fish require heavy metals (e.g. Fe, Mn, Cu, Zn, and Cr) within an acceptable trace amount to function and survive [1]. An excessive amount of Hg, As, Pb, and Cd elements could be detrimental to the living cells, and a prolonged exposure to the body can lead to illness or death [2]. Consequently, metals that were widely used to promote food production caused severe pollution worldwide, and thus had become a global concern in the past century [3–6]. Within the aquatic environment, such metals are considered as significant pollutants due to their intrinsic persistence, toxicities, non-biodegradable properties, and propensity to bioaccumulate up the food web [7–10]. At the end point, the intake of the metals will be a menace for anyone when heavy metals are consumed at a rate higher than the safe limit [11].

In the human body, alongside with a higher concentration, ineffective catharsis process can also make the heavy metals harmful even with a lower concentration [12]. For example, prostatic proliferative lesions, lung cancer, bone fractures, kidney failures are to be due to chronic exposure to Cd, even at a low concentration of ∼ 1 mg/kg [6, 13]. Cd is also deemed as a causative agent for long time exposure of skin, vascular, nervous system dysfunction, reproductive problems, and finally lead to cancer [14, 15]. Pb is termed as a non-essential element and can have detrimental health effect on human organ [16] including the nervous system, mental retardation, skeletal hematopoietic function disorder caused to even death [8]. Though Cr is crucial content for diet in terms of lipid metabolism and insulin activation [17], it can cause carcinogenic effects on human health [18, 19]. Cu plays essential role for enzyme functioning and hemoglobin synthesis [20, 21], however could also be a causative agent for toxicity accelerating nausea, bowel pain, diarrhea along with fever [22].

The aquatic organisms are directly or indirectly affected by these contaminants largely sourced from industrialization and urbanization [23–25]. Fish occupies higher trophic level in the food chain and are one of the most common bioindicators for pollutants [26, 27]. For many years, fishes are considered as a major protein supplier in human food consumption. Thus, the human body is largely susceptible to be enriched by a higher level of heavy metal concentration [28].

Generally, biomagnification occurs due to longstanding anthropogenic activities within a coastal ecosystem [29]. The accumulation of heavy metals in fish organs could also be driven by physiochemical and biological variables such as pH, temperature, hardness, exposure duration, feeding habits of species and habitat complexity [30]. While terrestrial species exhibit a strong pattern of biomagnification, marine and estuarine organisms show less clear pattern [31]. Condition factor, based on length-weight relationship, is one of the most common tools that is widely used in stock measurement model and to assess the life condition, reproduction records, health condition, and life cycle of a fish species [32]. Along with that, condition factor also suggest the food availability and quality, breeding duration, and process for distinct populations [1, 33]. In addition, this tool indicates the status of fish health due to stress in the population within an ecosystem [34].

The Karnaphuli river estuary is one of the potential fish population habitats along the southeast coast of Bay of Bengal known to be an important breeding, feeding and nursery ground for many aquatic species. At present, the ecosystem is receiving untreated effluents from several industries including textile crafts, dying industries, and others as it passes through the industrial zone [35]. A number of studies attempted to assess the contamination status from river and estuarine environment from Bangladesh [3, 36–39], from China [40], from Turkey [41]. However, to date there has been no proper investigation carried out on the potential human health risk evaluation due to heavy metal contamination in the fish species harvested and consumed from the Karnaphuli estuarine water body. The present study therefore aims to fill this knowledge gap by assessing the concentration of heavy metals in some selective fish species and their bioaccumulation status in relation to length-weight relationship and condition factor, and the human health risk evaluation for local adult and children consumers.

## Materials and methods

### Ethical statement

Live specimens from wild populations were collected from local fishermen. None of the sampled species were endangered or protected. No permit was required to conduct the study on invertebrates. There were no ethical considerations linked to the experiment.

### Sampling

The Karnaphuli River estuary was selected as the study area located from 22.234008 N and 91.821105 E to 22.289695 N and 91.794403 E (Fig. 1). A total of six commercial species (i.e. *A. bato, P. chinensis, L. parsia, M. cephalus, H. limbatus, and T. toil*) were collected from fishermen for a period of seven months (February 2018 to August 2018) using seine net. The collected samples were kept in plastic ice container and immediately stored in –12^0^ C. Afterwards in the lab, total length (cm) and weights (gm) were measured carefully to the nearest 0.1 cm using a vernier caliper; total weight was determined with an electronic balance to 0.01 g accuracy. Muscles of each specimen were dissected with stainless steel scissors. The dissected samples were then set for further chemical analysis using inductively coupled plasma mass spectrometry (ICP-MS, Model: ELAN9000, Perkin-Elmer, Germany) for metal detection. Data were analyzed statistically by fitting a straight line adopting the least square method.

**Figure 1.**
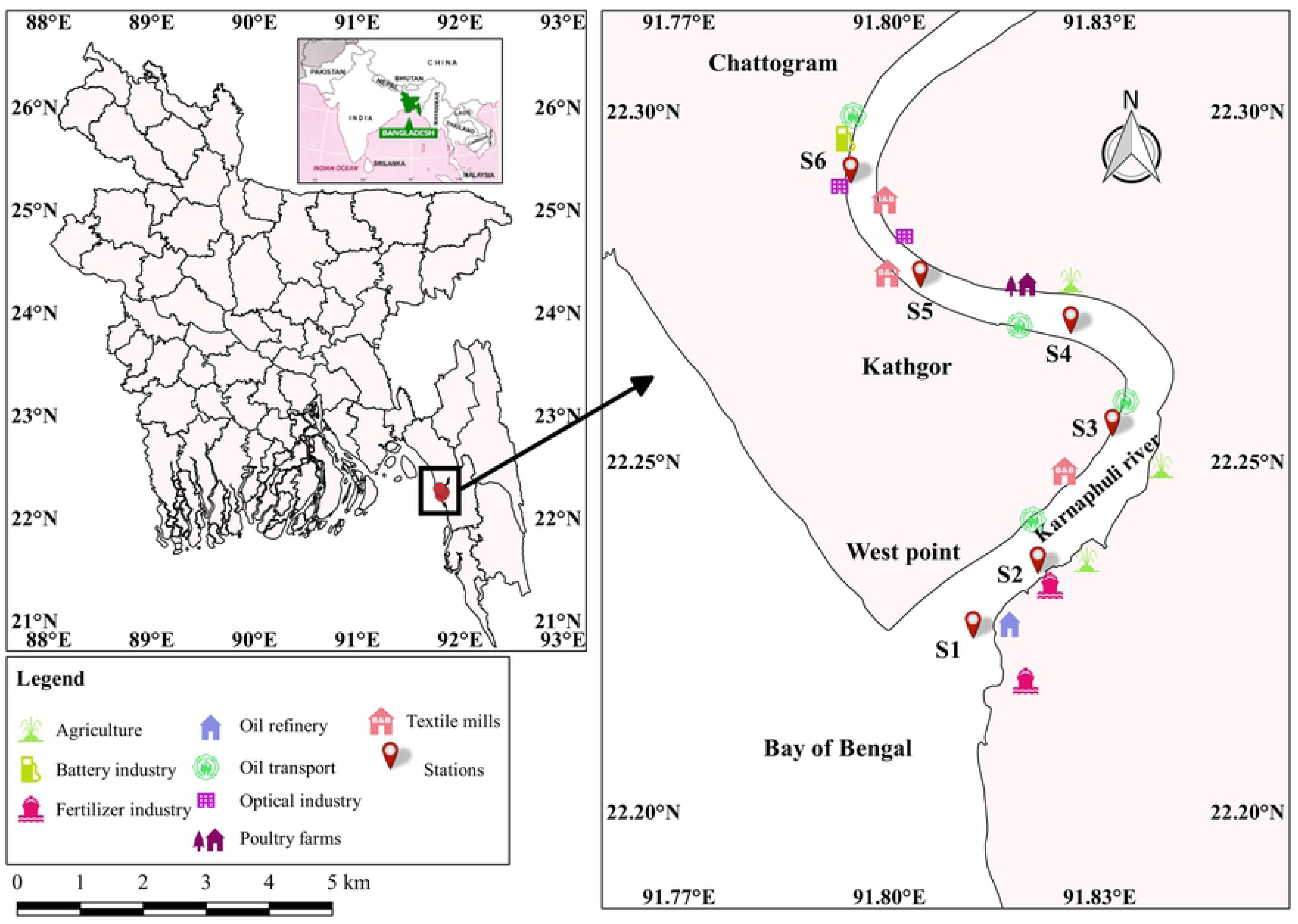
Lacation of the six sampling areas along with the source of introduction of the metal into the Karnaphuli river estuary.

### Chemical analysis procedures

A 1.5 g of dissected muscle portion was dried in an oven at 150^0^ C and then cooled. Afterwards, 3 ml of high concentrated H_2_SO_4_ and NHO_3_ was added and thoroughly mixed with the samples. The solution was heated on an oil bath adding ¾ drops of H_2_O_2_, repeated until the mixture was clear. The solution was mineralized using microwave digester (WX-6000, China). A standard reagent (Mark VI; Germany) was used to analyze the prepared samples in triplicate and the accuracy was obtained between 0–4% and 15%. Herein, the analytical accuracy was established at less than 10%.

### Metal pollution index (MPI)

To assess the metal pollution, the Metal pollution index (MPI) was adopted following [42] and [43]. The equation is as follows:

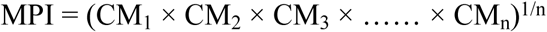

where, CM_1_ is the concentration value of first concerned metal, CM_2_ is the concentration value of second concerned metal, CM_3_ is the concentration of third concerned metal, CM_n_ is the concentration of n*_th_* metal (mg/kg dry wt) in the tissue sample of a certain species.

### Statistical analysis

The mean and standard deviation of metal concentrations were calculated. The Kolmogorov-Smirnov, Shapiro-Wilk and Kruskal-Wallis tests were performed using SPSS 23. Kolmogorov-Smirnov and Shapiro-Wilk tests were performed to identify the data dispersion avoiding the problems like normal/non-normal of data distribution in the aquatic ecosystem [44]. The Kruskal-Wallis test was carried out to identify significant variance of the targeted elements in the specimens of the studied area where p ≤ 0.05 was used as the cutoff for significance (confidence level in 95%). The employed correlation among the metals was classified in two groups [16], some correlations are positive and some were negative [45–48]. To identify the similar groups of the elements at the sampling sites, cluster analysis was executed with special variability [3, 49]. Resemble metals were in line in one cluster, while the dissimilar group of elements was plotted in another cluster to identify the term of contamination status [50–52].

### Bioaccumulation factor

Bioaccumulation factors (BAFs) were calculated as a ratio between the concentration level of biota (those in water) and the living environment of the specimens and was expressed as follows [53–55]:

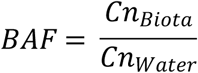

where *Cn_Biota_* is the concentration of metal in the tissues (mg/kg) and *Cn_Water_* is the metal concentration in the aquatic environment (mg/l). BAF is categorized as follows: BAF < 1000: no probability of accumulation; 1000 < BAF < 5000: bioaccumulative; BAF > 5000: extremely bioaccumulative [56].

### Length-weight relationship and condition factor

The length-weight relationship of the fish samples were calculated using Fulton condition factor following the equation [57–60]:

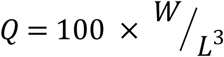

Where W is the total body weight of fish (gm), L is the total length of fish (cm). Fulton’s Q is categorized as follows: Q = 1: Condition is poor, Q = 1.2: condition is moderate, Q ≥ 1.40: condition is proportionally good [61]. The equation can be expressed by the following formula [1, 62]:

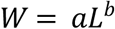

The equation can be estimated using the least-square formula adopted with the logarithm form of the equation is shown as [63]:

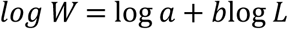

where, ‘a’ is the calculated intercept of the regression line, and ‘b’ is the coefficient of that regression. The ‘b’ values signify the growth pattern of an organism which can be classified as follows: b < 3: negative allometric, b = 3: Isometric and b > 3: positive allometric [57, 64].

As Fulton’s Q is substantially correlated with the length-weight relationship, exponent ‘b’ acts an identical role of determining the well-being of the organisms [57]. The deviation of the condition, further, depends on the food availability and the divergence of reproductive organ development [65].

## Human health risk

### Estimated daily intake (EDI)

Estimated daily intake (EDI) was calculated by the following equation [19, 66, 67]:

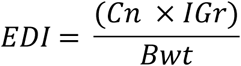

Where, C*n* is the concentration level of metal in the selected fish tissues (mg/kg dry-wt); *IGr* is the acceptable ingestion rate, which is 55.5 g/day for adults and 52.5 g/day for children [68, 69]; *Bwt* is the body weight: 70 kg for adults and 15 kg for children [68].

### Target hazard quotient (THQ) for non-carcinogenic risk assessment

THQ was estimated by the ratio of EDI and oral reference dose (RfD). RfDs of the different metals for example As, Pb, Cd, Cr, and Cu are 0.0003, 0.002, 0.001, 0.003 and 0.3, respectively [68, 70]. The value of ratio < 1 implies a non-significant risk effects [71]. The THQ formula is expressed as follows [69, 72–74]:

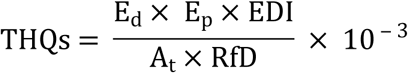

Where *E_d_* is exposure duration (65 years) (USEPA, 2008); *E_p_* is exposure frequency (365 days/year) [18]; *A_t_* is the average time for the non-carcinogenic element (*E_d_*×*E_p_*).

### Hazard index (HI)

Hazard index (HI) was calculated for the multiple elements (Hg, As, Mn, and Cr) found in the fish samples and the equation is as follows [3, 16, 75, 76]:

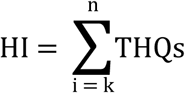

where, THQs is the estimated risk value for individual metal [77]. When HI value is higher than 10, the non-carcinogenic risk effect is considered high for exposed consumers [78–80].

### Carcinogenic risk (CR)

To assess the probability of developing cancer over a lifetime, the carcinogenic risk is evaluated for the consequence of exposure to the substantial carcinogens [81, 82]. The acceptable range of the risk limit is 10^−6^ to 10^−4^ [83–86]. CRs higher than 10^−4^ are likely to increase the probability of carcinogenic risk effect [87, 88]. The established equation to assess the CR is as follows [69, 70, 89, 90]:

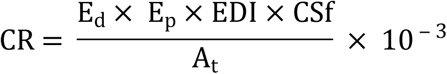

Where CSf is oral slope factor of particular carcinogen (mg/kg-day) [83]. Available CSf values (mg/kg-day) are: As (1.5), Pb (0.0085) and Cd (6.3) [83].

## Results

### The concentration of heavy metals and source identification

The average concentration of Pb, Cu, As, Cr, and Cd from the fish tissues were 13.88 (range: 3.19–6.19); 12.10 (range: 10.27–16.41); 4.89 (range: 3.19–6.19); 3.36 (range: 2.46 – 4.17), and 0.39 (range: 0.21–0.74) respectively. The maximum mean concentration was Pb and minimum was Cd (please see Table 1).

**Table 1.** Concentration of heavy metals of different species and their feeding nature, length and weight and a comparison of other relevant studies along with various standard guideline values.

| Species | Feeding nature | Amounts | Length (cm) | Weight (gm) | Heavy metals (mean ± std) mg/kg | | | | | MPI | References |
| --- | --- | --- | --- | --- | --- | --- | --- | --- | --- | --- | --- |
|  |  |  |  |  | As | Pb | Cd | Cr | Cu |  |  |
| *A. bato* | Demersal | 18 | 13.26±1.35 | 72.64±6.39 | 4.65±1.06 | 15.22±1.32 | 0.60±0.13 | 3.54±0.54 | 15.29±3.82 | 4.70 | |
| *P. chinensis* | Demersal | 18 | 6.96±0.90 | 166.58±21.68 | 5.03±0.86 | 14±1.79 | 0.44±0.14 | 3.59±0.55 | 13.10±2.49 | 4.29 | |
| *L. parsia* | Demersal | 18 | 7.34±1.17 | 25.20±3.12 | 4.36±0.93 | 13.98±1.93 | 0.34±0.09 | 3.30±0.40 | 9.50±1.16 | 3.65 | |
| *M. cephalus* | Pelagic | 18 | 27.17±2.30 | 711.44±111.03 | 4.89±0.48 | 12.70±1.72 | 0.31±0.09 | 3.14±0.36 | 11.48±1.27 | 3.69 | |
| *H. limbatus* | Pelagic | 18 | 10.97±0.80 | 26.77±4.39 | 5.14±0.86 | 13.77±1.54 | 0.35±0.08 | 3.52±0.48 | 12.52±1.40 | 4.05 | |
| *T. toli* | Pelagic | 18 | 30.68±2.54 | 646.82±41.16 | 5.26±0.49 | 13.61±0.82 | 0.31±0.07 | 3.11±0.54 | 10.72±1.47 | 3.75 | |
| **Water(ml/l)** | | | | | 0.006±0.003 | 0.017±0.006 | 0.002±0.001 | 0.006±0.002 | 0.006±0.001 | | |
| **Guidelines** |  |  |  |  |  |  |  |  |  |  |  |
| FAO |  |  |  |  | 1 | 2.5 | 0.2 | 1 | 10 |  | FAO, 1983 |
| WHO |  |  |  |  | 0.01 | 2 | - | 0.15 | 3 |  | WHO, (1985) |
| EU |  |  |  |  | - | 0.1 | 0.05 | 1 | 3 |  | EU, (2001) |
| Bangladesh (fish) |  |  |  |  | 5 | 0.30 | 0.25 | - | 5.00 |  | MOFL (2014) |
| **Literature** |  |  |  |  |  |  |  |  |  |  |  |
| Coastal area, Bangladesh |  |  |  |  | 0.08-13 | 0.07–0.63 | 0.03–0.09 | 0.15–2.2 | 1.3-14 |  | Raknuzzaman et al., 2016 |
| Arasalar River, India |  |  |  |  | - | 0.23 | 6.13 | 0.3 | - |  | Lakshmanasenthil et al., 2013 |
| North east coast, India |  |  |  |  | 0.64 | - | 0.33 | - | 3.9 |  | Kumar et al., 2012 |
| Ganga River, India |  |  |  |  | - | 3-6 | 0.1-2.9 | - | 10-100 |  | Mitra et al., 2012 |
| Pearl River, China |  |  |  |  | - | 8.64 | 8.55 | 8.73 | 2.48 |  | Kwok et al., 2014 |
| Meiliang Bay, China |  |  |  |  | - | 0.636 | 0.173 | 0.118 | 0.336 |  | Rajeshkumar et al., 2018 |
| Iskenderun Bay, Turkey |  |  |  |  | - | 0.09–6.95 | 0.01–4.16 | 0.07–6.46 | 0.04–5.43 |  | Türkmen et al., 2005 |

The evaluated MPIs ranged from 3.65 mg/kg to 4.70 mg/kg with the mean of 4.02 mg/kg (Table 1). Due to higher concentration level, the maximum MPI value (4.70 mg/kg) was corresponded to *A. bato, followed by P. chinensis* (4.2 mg/kg) *H. limbatus* (4.05 mg/kg), *T. toli* (3.75 mg/kg), *M. cephalus* (3.69 mg/kg), and *L. parsia* (3.65 mg/kg).

Kolmogorov-Smirnov and Shapiro-Walk test revealed that the metals in the targeted fish species were non-normally distributed along the study area. The adopted Levene tests adopted and resulted that metals were non-homogenouosly distributed. Kruskal-Wallis test identified that the distribution of metal was significantly different (p ≤ 0.05) in the fish species along the sampling stations. From Table 2, among the species, *L. parsia and H. limbatus* exhibited significant relationship (regression line) with As, while *As, T. toil* was with As and Pb. None of the other metals showed a significant linear relationship with the organisms. Among the species, *T. toil* showed the maximum response for Pb (R^2^=99.5%), followed by *L. parsia* for As (R^2^=83.7%).

**Table 2:** A regression analysis between metal distributions in the selected specimens.

| Species and metals | Regression equation | Standard error (b) | Pearson's r | P values | R<sup>2</sup> values (%) |
| --- | --- | --- | --- | --- | --- |
| *A. bato* |  |  |  |  |  |
| As | $y = 8.66 + 0.99x$ | 0.404 | 0.774 | 0.070 | 58.1 |
| Pb | $y = 11.88 + 0.09x$ | 0.511 | 0.088 | 0.867 | 0.7 |
| Cd | $y = 13.44 - 0.31x$ | 5.166 | -0.029 | 0.955 | 0.01 |
| Cr | $y = 12.70 + 0.16x$ | 1.25 | 0.063 | 0.904 | 0.4 |
| Cu | $y = 11.56 + 0.11x$ | 0.168 | 0.313 | 0.545 | 7.5 |
| *P. chinensis* |  |  |  |  |  |
| As | $y = 4.19 + 0.55x$ | 0.449 | 0.522 | 0.287 | 13.3 |
| Pb | $y = 5.14 + 0.13x$ | 0.243 | 0.257 | 0.621 | 2.6 |
| Cd | $y = 5.72 + 2.81x$ | 2.993 | 0.425 | 0.4 | 5.8 |
| Cr | $y = 5.75 + 0.34x$ | 0.8 | 0.205 | 0.695 | 4.2 |
| Cu | $y = 4.58 + 0.18x$ | 0.157 | 0.502 | 0.309 | 25.3 |
| *L. parsia* |  |  |  |  |  |
| As | $y = -0.96 + 0.73x$ | 0.16 | 0.914 | **0.01** | 83.7 |
| Pb | $y = 7.79 + 0.84x$ | 0.711 | 0.51 | 0.301 | 26.0 |
| Cd | $y = 0.12 + 0.03x$ | 0.036 | 0.378 | 0.459 | 14.3 |
| Cr | $y = 1.86 + 0.2x$ | 0.139 | 0.580 | 0.227 | 0.1 |
| Cu | $y = 4.24 + 0.72x$ | 0.346 | 0.719 | 0.107 | 51.7 |
| *M. cephalus* |  |  |  |  |  |
| As | $y = 2.08 + 0.11x$ | 0.09 | 0.5 | 0.311 | 25.0 |
| Pb | $y = 23.98 - 0.42x$ | 0.309 | -0.557 | -0.557 | 31.0 |
| Cd | $y = 0.87 - 0.02x$ | 0.016 | -0.547 | 0.26 | 30.0 |
| Cr | $y = 4.04 - 0.03x$ | 0.075 | -0.216 | 0.68 | 4.7 |
| Cu | $y = 16.95 - 0.21x$ | 0.257 | -0.364 | 0.477 | 13.3 |
| *H. limbatus* |  |  |  |  |  |
| As | $y = -4.63 + 0.89x$ | 0.309 | 0.821 | **0.045** | 67.5 |
| Pb | $y = 1.09 + 1.156x$ | 0.778 | 0.596 | 0.211 | 35.6 |
| Cd | $y = 0.36 - 0.01x$ | 0.048 | -0.014 | 0.978 | 0.01 |
| Cr | $y = -0.6 + 0.38x$ | 0.238 | 0.620 | 0.189 | 38.5 |
| Cu | $y = 2.54 + 0.91x$ | 0.749 | 0.519 | 0.291 | 26.9 |
| *T. toli* |  |  |  |  |  |
| As | $y = 0.48 + 0.16x$ | 0.059 | 0.799 | **0.05** | 63.9 |
| Pb | $y = 3.74 + 0.32x$ | 0.012 | 0.997 | **1.11E-5** | 99.5 |
| Cd | $y = 0.35 - 0.01x$ | 0.013 | -0.042 | 0.936 | 0.2 |
| Cr | $y = -1.22 + 0.14x$ | 0.08 | 0.661 | 0.152 | 43.8 |
| Cu | $y = 0.37 + 0.34x$ | 0.236 | 0.580 | 0.227 | 33.7 |

The Pearson correlation among the metals in different species was presented in Table 3. In our study, Cd and Pb were found significantly correlated with each other (r = 0.88). Also, Cr was significantly and positively correlated with Pb (0.664) and Cd (0.698). Cu also showed a significant positive association with Cr (r = 0.704).

**Table 3.**
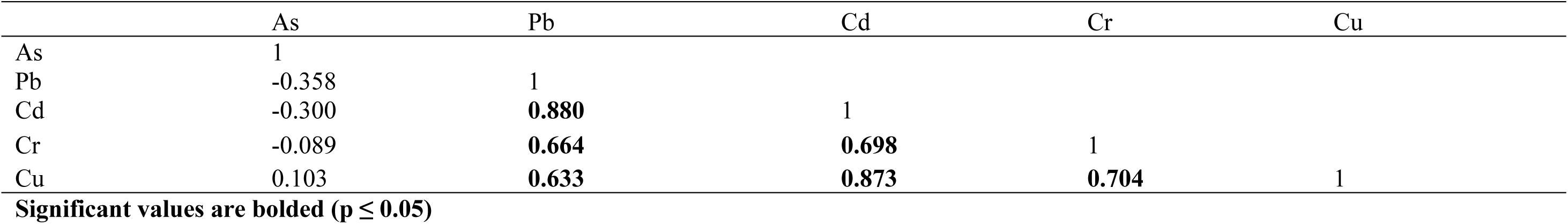
Pearson’s correlation matrix among the metals.

**Table 4:** Length-weight relationship, growth pattern and Fulton condition factor for the targeted fishes

| Species | a (intercept) ± SE | b (slope regression) ± SE | Group (Growth pattern) | $W = aL^b$ | Fulton's Q ± std | Fish Condition |
| --- | --- | --- | --- | --- | --- | --- |
| *A. bato* | 19.62 ± 16.63 | 3.99 ± 1.24 | Positive allometric | $W = 19.62 \times L^{3.99}$ | 3.22 ± 0.78 | Good |
| *P. chinensis* | 7.23 ± 25.94 | 22.88 ± 3.7 | Positive allometric | $W = 7.23 \times L^{22.88}$ | 51.77 ± 15.26 | Good |
| *L. parsia* | 7.13 ± 3.8 | 2.76 ± 0.51 | Isometric | $W = 7.13 \times L^{2.46}$ | 7.10 ± 3.44 | Good |
| *M. cephalus* | 0.002 ± 0.0008 | 10.85 ± 4.98 | Positive allometric | $W = 0.002 \times L^{10.85}$ | 3.56 ± 0.45 | Good |
| *H. limbatus* | 0.01 ± 0.002 | 2.43 ± 0.41 | Negative | $W = 0.01 \times L^{2.43}$ | 2.03 ± 0.23 | Good |
| *T. toli* | 164.30 ± 60.97 | 15.72 ± 1.98 | Positive allometric | $W = 164 \times L^{15.72}$ | 2.29 ± 0.46 | Good |

In the present study, Ward-Linkage method was employed with Euclidean distance, which resulted in three distinct clusters, presented in Fig 2. Cluster 1 included As, whereas, Pb, Cd, and Cu confined in cluster 2, whereas Cr was found in cluster 3.

**Figure 2.**
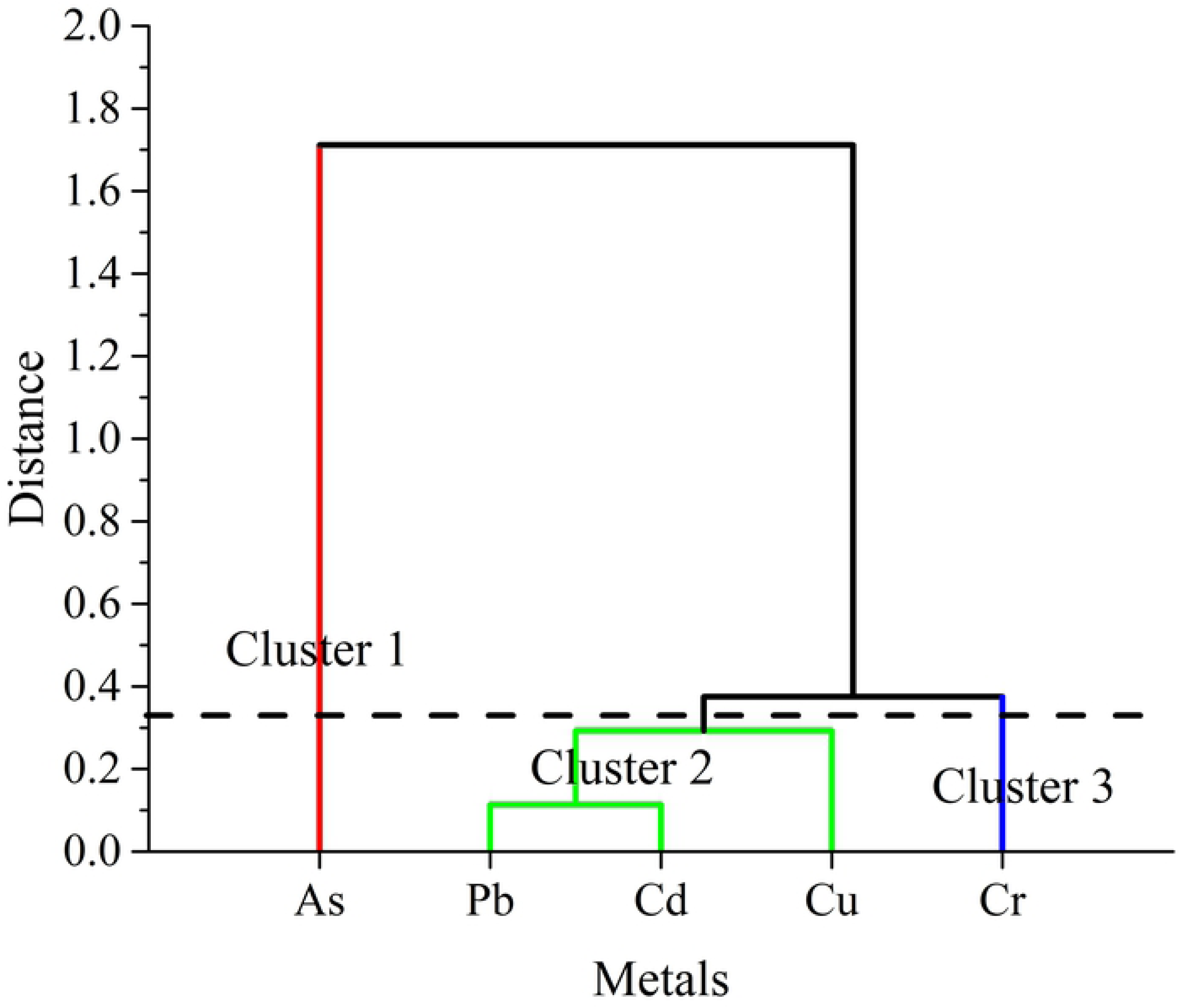
Hierarchical cluster (dendrogram) using Ward linkage method among the experimented metals in fish species.

### Length-weight Relationship and condition factor evaluation

Higher ‘b’ value reflected the appetite state and reproductive organ development of the species [64]. The identified b value of *L. parsia* from the length weight relatioship was close to 3, hence, it represented isometric growth pattern that was considered as ideal shape. Meanwhile, among all, species, *P. chinensis* exhibited the highest positive allometric growth, which was 37 times higher, than average value and 14 folds, on average, higher than other species.

### Bioaccumulation (BAF) status of targeted species

The estimated BAFs were depicted in Fig 3. The BAFs were ranged from 110.53 for Cd observed in S4 to 3353.7 for Cu as well. The minimum value was found for *Tenualosa toil*, on the other hand, *Apocryptes bato* showed maximum bioaccumulation result. Moreover, the mean BAFs of the metals were observed in the species as follows: Cu (1971.42) > As (1042.93) > Pb (913.66) > Cr (864.99) > Cd (252.03).

**Figure 3.**
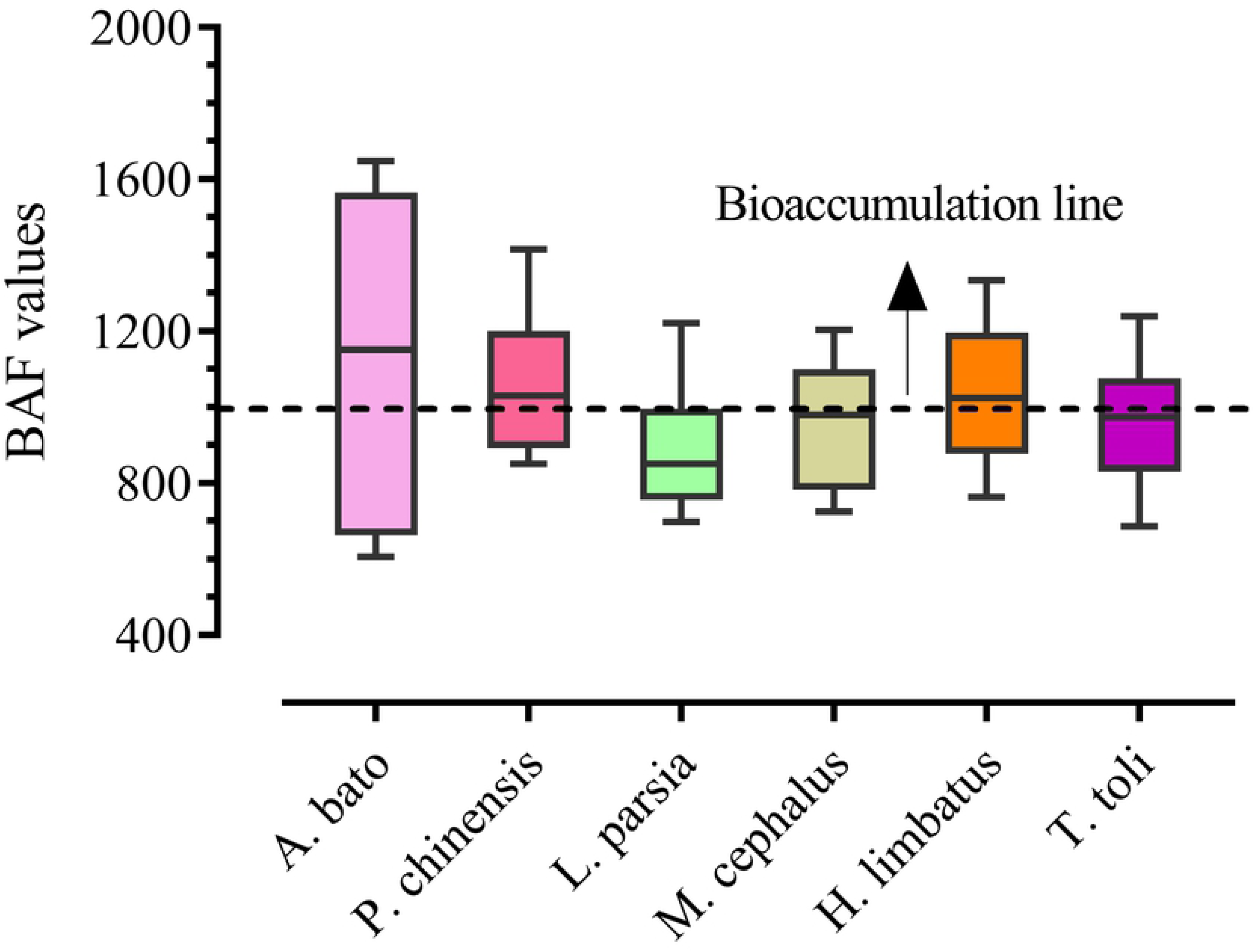
Bioaccumulation factor among the species that were varied from particular metals and species.

## Health risk evaluation

### Estimated daily intake (EDI)

The explored EDI of two concerned age groups, adults and children, was presented and summarized in Table 5. The study noted that adults and children showed comparably higher EDIs for demersal species than pelagic ones. In the consequence, high doses of demersal species were exposed to the consumers through consuming metal affected fish species as the food items. The EDIs for both groups were organized in the following order: Pb > Cu > As > Cr > Cd.

**Table 5:** A comparison between recommended daily allowance (RDA) and estimated daily intake (EDI) for adults and children.

| Elements | Mean concentration<br>(mg/kg) | RDA (mg/kg/person) * | EDIs (mg/day/person) |  |
| --- | --- | --- | --- | --- |
|  |  |  | Ad | Ch |
| As | 4.89 | 0.15 | 0.005 | 0.029 |
| Pb | 13.88 | 0.25 | 0.011 | 0.049 |
| Cd | 0.39 | 0.07 | 0.001 | 0.001 |
| Cr | 3.36 | 0.23 | 0.003 | 0.012 |
| Cu | 12.10 | 35 | 0.010 | 0.042 |

\*WHO, 2000

### Target hazard quotient (THQ) for non-carcinogenic risk

The assessed target hazard quotient (THQ) for the studied fish species were displayed in Table 6. THQs from the study area in the adult group induced to As, Pb, Cd, Cr, and Cu were 0.016, 0.003, 3.0E-04, 0.001 and 2.38E-04, respectively, where for children were 0.097, 0.014, 0.001, 0.004 and 0.001, respectively. Moreover, the rank of the THQs of the elements was as follows: As > Pb > Cr > Cd > Cu. While, for the cumulative scenario of HI, children were 5.83 times more susceptible than adults. However, the investigated HI was not surpass the recommended limit (Table 6).

**Table 6.** Calculated THQ, HI and CR for the selected two aged groups.

| Species | THQ (As) |  | THQ (Pb) |  | THQ (Cd) |  | THQ (Cr) |  | THQ (Cu) |  | HI |  | CR (As) |  | CR (Pb) |  | CR (Cd) |  |
| --- | --- | --- | --- | --- | --- | --- | --- | --- | --- | --- | --- | --- | --- | --- | --- | --- | --- | --- |
|  | Ad | Ch | Ad | Ch | Ad | Ch | Ad | Ch | Ad | Ch | Ad | Ch | Ad | Ch | Ad | Ch | Ad | Ch |
| *A. bato* | 0.012 | 0.054 | 0.003 | 0.015 | 0.000 | 0.002 | 0.001 | 0.004 | 0.000 | 0.001 | 0.017 | 0.077 | 5.47E-06 | 2.44E-05 | 1.79E-05 | 8.00E-05 | 2.99E-06 | 1.30E-05 |
| *P. chinensis* | 0.013 | 0.059 | 0.003 | 0.014 | 0.000 | 0.002 | 0.001 | 0.004 | 0.000 | 0.001 | 0.018 | 0.08 | 5.93E-06 | 2.64E-05 | 9.35E-08 | 7.40E-05 | 2.18E-06 | 9.70E-06 |
| *L. parsia* | 0.011 | 0.051 | 0.003 | 0.014 | 0.000 | 0.001 | 0.001 | 0.004 | 0.000 | 0.001 | 0.016 | 0.071 | 5.14E-06 | 2.29E-05 | 9.33E-08 | 4.20E-07 | 1.68E-06 | 7.50E-06 |
| *M. cephalus* | 0.013 | 0.057 | 0.003 | 0.013 | 0.000 | 0.001 | 0.001 | 0.004 | 0.000 | 0.001 | 0.017 | 0.076 | 5.77E-06 | 2.57E-05 | 8.48E-08 | 3.80E-07 | 1.52E-06 | 6.80E-06 |
| *H. limbatus* | 0.013 | 0.06 | 0.003 | 0.014 | 0.000 | 0.001 | 0.001 | 0.004 | 0.000 | 0.001 | 0.018 | 0.08 | 6.06E-06 | 2.7E-05 | 9.2E-08 | 4.10E-07 | 1.72E-06 | 7.70E-06 |
| *T. toli* | 0.030 | 0.3 | 0.003 | 0.014 | 0.000 | 0.001 | 0.001 | 0.004 | 0.000 | 0.001 | 0.034 | 0.319 | 1.35E-05 | 1.35E-04 | 9.09E-08 | 4.00E-07 | 1.54E-06 | 6.90E-06 |
| Mean | 0.016 | 0.097 | 0.003 | 0.014 | 0.000 | 0.001 | 0.001 | 0.004 | 0.000 | 0.001 | 0.015 | 0.079 | 6.98E-06 | 4.36E-05 | 3.07E-06 | 2.58E-05 | 1.94E-06 | 8.64E-06 |

### Carcinogenic risk (CR)

Exposure of CR was estimated for a particular element and summarized in Table 6. The measured CR values of As, Pb and Cd were ranged from 5.14E-06-1.35E-05, 8.48E-08-1.79E-05 and 1.52E-06-2.99E-06 respectively in adults and 2.29E-05-1.35E-04, 3.78E-07-7.99E-05 and 6.76E-06-1.33E-05 in children. The result showed that children were exposed to higher CRs than adults. But, calculated CR values for both aged groups were noted far from the risk acceptable range (10^−6^ to 10^−4^).

## Discussion

### The concentration of heavy metals and source identification

In the present study area, *T. toil* showed the highest concentration of As, whereas *A. bato* exhibited the maximum concentration for Pb, Cd, and Cu, and metal, Cr was belonged to at the highest concentration in *P. chinensis.* The concentrations of the metal in the editable tissues were ranked in the following order: *A. bato > P. chinensis > H. limbatus > T. toli > M. cephalus > L. parsia.* In terms of MPI, the fact is that, the demersal organisms are closely related to the sediment that is to be indirect and long-term source for contamination assessment [91]. Present study showed that, demersal species had comparatively higher concentration level of metals than pelagic ones. The value of MPI for the present study were ranged from 3.65 to 4.70, whereas MPI values for Tilapia, *Sarotherodon melanotheron* and Silver catfish, *Chrysichthys nigrodigitatus* were found much higher (8.1 to 17.76) from Okrika Estuary, Nigeria. This is most likely due to the oil bunkering and transportation activities along the study sites [92]. The findings of MPI of the present study were almost similar to that of *Rutilus rutilus* in Pluszne Lake [1]. The metal accumulation in fishes could be highly influenced by sampling locations and habitats [93, 94].

In terms of source identification, industrial operation and antifungal wood preservative frequently use As in production, which could deteriorate the water and sediment quality [95]. In the study area, there are several manufactures industries largely use alloy, sheep, leather technologies, paints, poisonous chemicals contains As. Modern day microelectronic and optical industries use heavy metals for their commercial aspect which is termed as notable sources for As intrusion in the aquatic environment [96]. Nonessential element, Pb comes from extreme agriculture, poultry forms, industries, and textile mills to the aquatic ecosystem [97] which was the source of the metal in the study area. Thus the benthic feeders are to be greatly affected by the deposited Pb in the ecosystem [22]. Cd metal was typically found at a low concentration in the aquatic environment, however, incognizant use of phosphate fertilizer and industries are two primary sources of Cd introduction [98]. In the study area, largely operated nickel-cadmium battery manufactories along with industries engaged with Cd metal incineration and production may increase the Cd concentration level in the aquatic environment [99, 100]. Beside Cd and Pb, Cr is also widely introduced in textile industries [101]. Near to the bank of the Karnaphuli estuary, such commercial textile industries produce color pigment and thus become a common contaminant for the aquatic ecosystem [102]. Notably, a considerable level of Cu become swelled up in the study area due to oil droppings from ships and boats, recurrent usage of antifouling paints and other boating interferes [20].

The concentration level of As and Pb from our study was higher than all other findings and recommended guidelines. Although Cd concentration was at lower level comparing Northeast coast, Ganga river, and Pearl river, it surpassed the all guideline values along with other coastal environments, Meiliang Bay, Iskenderun Bay, Arasalar river and Coastal area of Bangladesh. Similar findings were observed for Cr that crossed the recommended limit and other comparable studies except for Pearl river. On the other hand, the Ganga river showed a higher concentration for Cu, while specimens showed almost 4 times deviation from the accepted limits except for *L. parsia*.

### Bioaccumulation (BAF) status of targeted species

Bioaccumulation potential of metals was assessed in muscles of various fish species, which were varied from species to species. The hierarchy suggest that most of the species were tend to be bioaccumulative as the value approaches near to 1000. *A. bato* exhibited the highest concentration of bioaccumulation in the studied area. The accumulation of the metal elements in an aquatic organism depends upon the classification of species, invasion pathways, metabolic characters of the sampled tissues and finally, the surrounding the environmental status of the species living in [103]. In our result, it was observed that, the BAFs of As, Cd, Cr, Pb, and Cu were relatively higher than that of Pearl river estuary [104], where the BAFs in tilapia were reported in the following order: Cd > Cu > Pb > Cr. Such reports were mostly in line with our results. The fact is that, Cu is left persisted actively in muscles due to being an essential element of living tissue [69, 105]. Notably, the capability of bioaccumulation of Cd takes a long time to spare that makes it relatively infirm [8].

### Human Health risk evaluation

EDI, based on the oral reference dose (RfD) for an individual element [106] reflects the daily exposure to the toxic element and is executed to avoid any harmful effect on human health [72]. The records of EDI of the people were compared with recommended daily allowance (RDA), provided by WHO, and introduced that, mean EDI values of the metals were still lower than RDAs. Values, which were lower than RDA guidelines, revealed a lower possible health effect to the consumers for those elements. But, it would be unwise to take it as a permanent measurement to reach a final conclusion describing as ‘acceptable value’ and ‘unacceptable value’ when the doses were lower than RDAs or Rfds [70, 72].

The value found for THQs for both adult and childen were below 1, revealing that adverse effect on human health might not occur. Similarly, HI result also followed the THQ trend. Hence, there is no such potential non-carcinogenic effect for the consumers due to intake of the fish species. Studies carried out by several authors in similar condition were in line with our results [43, 78, 107–109]. In general, the assessment of THQ for human health risk evaluation has no dose-response relation of the examined elements [110]. However, human can be suffered in the long run dramatically by the multiple pollutants simultaneously [86].

The CRs value lower than 10^−4^ indicated a negligible health risk. The CRs value found in this study suggested an acceptable limit and therefore, consumers are less prone to carcinogenic. In fact, 90% of the carcinogenic risk is observed for the As contaminated aquatic food items. The inorganic state of As is lethal than organic one [8, 111] and only 10% of total As can be assessed as inorganic form [72]. Present study was compared to [112] which found the carcinogenic risk value was in acceptable range (10^−6^ to 10^−4^), except for the metal of Cr which surpassed the CR limits. The reason was the rapid increase of metal pollution for the last 10 years, that phenomena support the study area. Again, in the Persian Gulf, consumers were at threshold limit for As in concern for carcinogenic risk [113]. For this reason, carcinogenic risk should be given more attention due to intake of aquatic products, especially for the study area.

## Conclusions

In the present study, the high concentration of metal was observed in *A. bato* for relatively high concentration level of Cu. Moreover, in most cases, metals concentration exceeded the recommended guideline limits. The maximum metal accumulation was recorded *A. bato* and species *A. bato, P. chinensis, H. limbatus* were observed as extreme bio-accumulative species in cumulative aspects. Lastly, CR assessment was also in acceptable threshold indicating that, local consumers were free from the sabotage of cancer risk for the time being but they might affected in future if still they consume fish from studied region. Finally, children were more vulnerable for health risk than adults. Nonetheless, further study required ensuring the same conclusions are reached.

## Acknowledgments

The authors acknowledged to the Rapid Action Battalion Headquarter, Bangladesh authority for providing necessary fund for heavy metal analysis and instrumental facilities with conventional techniques. Authors also would like to thank anonymous reviewers for improvement of this manuscript.

